# Mobile tuberigen impacts tuber onset synchronization and canopy senescence timing in potato

**DOI:** 10.1101/2023.11.08.566204

**Authors:** Bas van den Herik, Sara Bergonzi, Christian W.B. Bachem, Kirsten H. ten Tusscher

## Abstract

Yield of harvestable organs is a complex function of photosynthetic output, and sink-strength and timing of competing carbon sinks. In potato (*Solanum tuberosum*) the effect of tuber onset timing and post-tuberization canopy senescence on growth dynamics and tuber fresh weight are poorly understood. To advance our understanding we compared above- and belowground traits of wildtype plants (WT) with *StSP6A*, i.e., tuberigen, knockdown plants (SP6Ai) and developed simple computational models to aid interpretation of results. We find that SP6Ai results in a delay of approximately 2 weeks in tuber onset, yet has a 4-to-5-week delayed canopy senescence. Together this results in a prolonged tuber growth phase, with reduced synchronization in tuber onset and a resulting increased variance in tuber sizes, while overall final tuber fresh weight remains similar. Using a leaf and tuber growth model comparing various leaf senescence mechanisms, we find that resource competition, and not a shared signal for tuberization and senescence, is able to explain how delayed tuberization leads to further delayed senescence. Our results point to a role for resource competition in the correlated timing of tuber onset and canopy senescence, as well as a leading role for *StSP6A* in tuber onset synchronization and tuber size uniformity.

**Graphical Abstract:** 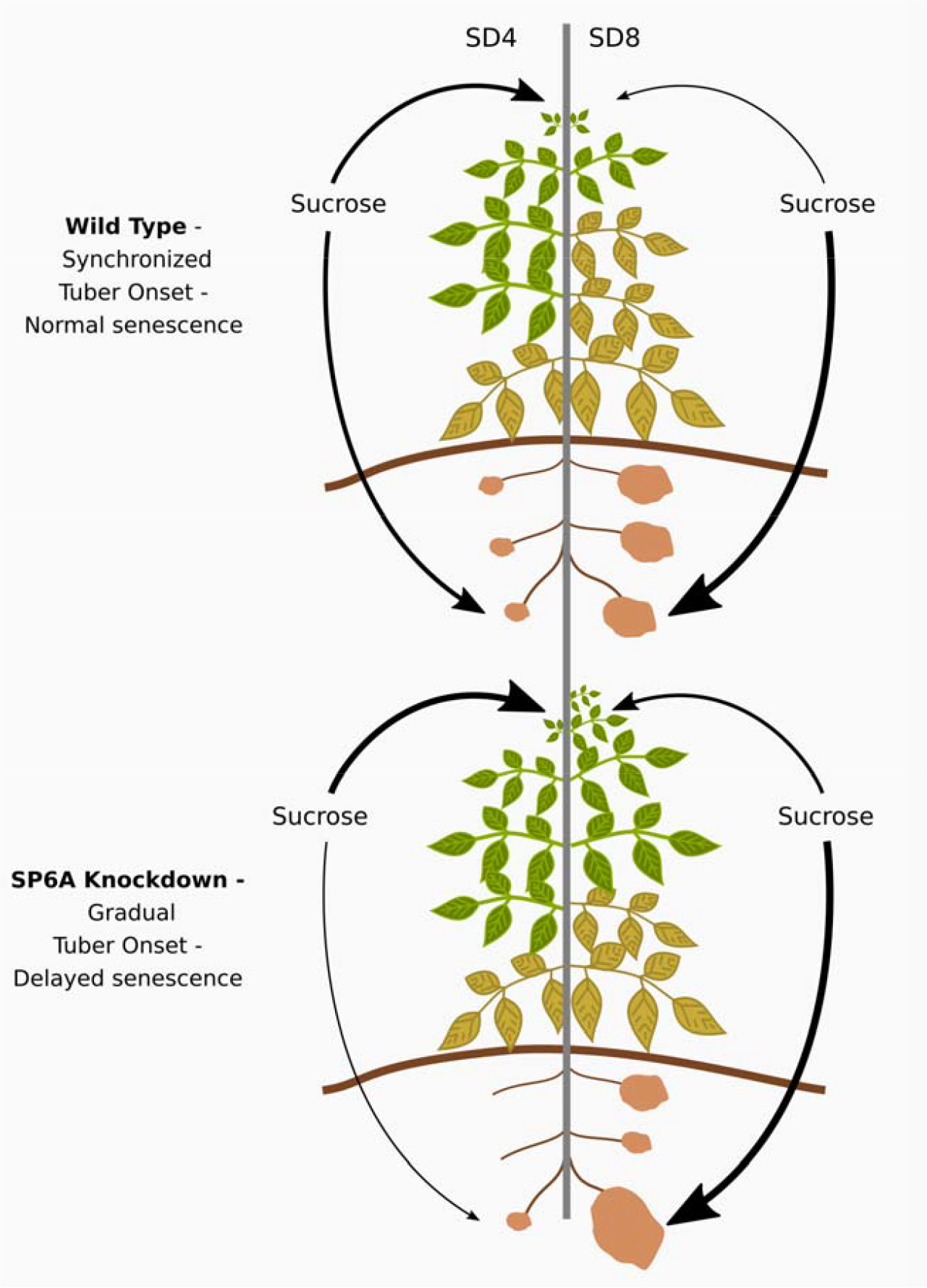

## 1. Introduction

Photosynthetic output and the subsequent distribution of produced assimilates across different sink organs play a vital role in determining the yield of harvestable plant organs. As such, both the strength and the timing of competition for sucrose between sink organs as well as the dynamics of source organs have a major impact on final crop yield. In potato, upon tuber formation, a new strong sink is formed coinciding with a switch from symplastic to apoplastic unloading (Viola et al., 2001). As a result, sucrose delivery to tubers is substantially increased at the cost of other plant organs (Fernie et al., 2020). *StSP6A*, the tuberigen, affects tuberization and sucrose delivery through initiation of tuber formation (Navarro et al., 2011) as well as increasing the relative efficiency of resource allocation towards tubers by mitigation of sucrose efflux via the sucrose exporter *StSWEET11* in the long-distance phloem (Abelenda et al., 2019; van den Herik et al., 2021; van den Herik & ten Tusscher, 2022). Tuberization is further under control of environmental conditions (such as light, temperature, water availability and soil nutrient levels), hormone levels and sucrose availability. For example, increased sucrose export from leaves leads to early flowering as well as increased tuber size (Chincinska et al., 2008). Interestingly, tuberization also occurs in *StSP6A* knockdown plants (Navarro et al., 2011), suggesting that other known tuberization signals such as the *StBEL5* factor, giberellin signalling and sucrose play partly redundant roles (Khondare et al., 2020). Indeed, recently it was shown that the florigen *StSP3D* also acts as a mobile tuber-inducing protein in the absence of the main tuberigen *StSP6A (*Jing et al., 2023). Thus, while developmental signals control the development of tubers through multiple, partly redundant, pathways, competition for and dependence on a shared energy resource leads to complex feedback relationships between timing and formation of tubers, leaves and other organs.

Overall canopy senescence timing (i.e., time point of decline of total leaf area) appears to parallel tuber development, with early tuber forming genotypes also undergoing earlier senescence. This phenomenon has been generally described as post-tuberization senescence. Leaf senescence is a natural developmental transition, which is dependent on organ age, as well as leaf nitrate and sucrose levels. Low nitrate, and both high and low sucrose levels have been described to promote leaf senescence (Rankenberg et al., 2021; Wingler, 2018). Thus, while tuber sink strength is enhanced after tuber formation, subsequent leaf senescence causes overall source-strength to diminish. An open question remains whether causal relations exist between these developmental events and how their precise relative timing impacts tuber yield. In addition to a competition for carbon between distinct organ types, (i.e. tubers, leaves, and flowers), tubers compete with each other for sucrose and this is likely an important factor determining potato tuber size distribution. Again, competition outcome depends on differences in sink strength, timing but also plant architecture, i.e. whether different sinks have different distances to sources (van den Herik & ten Tusscher, 2022; Zhu et al., 2021; Pallas et al., 2010). To what extent differences in tuber onset and growth timing merely reflect differences in prior stolon development, and to what extent regulation at the stage of tuberization onset may affect tuber size differences has not been systematically investigated. Open questions to address are thus 1) to what extent tuber development is controlled through competition for sucrose, 2) whether and how differences in tuber developmental timing affect agronomically important parameters such as tuber size distribution, and 3) whether post-tuberization canopy senescence is controlled largely in parallel with or is partly a downstream effect of tuber onset.

While previous studies have quantified growth of tubers and sugar/starch levels during development in field level experiments (Plaisted, 1957), during early tuberization (Davies, 1984) or tracked individual tuber volume development (Struik et al., 1988), there are no quantitative, plant level dynamic data sets to answer the questions above. Therefore, we set up an experiment aimed to generate a detailed data set covering the temporal development of individual potato plants, measuring various aspects of shoot and tuber growth, including sugar dynamics. We performed a climate chamber experiment under controlled day-length conditions with an obligate short-day tuber onset genotype, in WT and *StSP6A* RNAi (SP6Ai) lines. The experiment was set up to investigate resource competition between tubers and leaves, with the *StSP6A* RNAi lines expected to have delayed and possibly reduced tuberization. To aid the interpretation of experimental results, we combined our experiments with several simple computational models.

## 2. Materials & Methods

### 2.1 Experimental set-up and measured traits

#### 2.1.1 Plant material and growing conditions

Wild-type (*S. tuberosum* group Andigenum 7540 (referred to as S. *Andigena*)) and two *StSP6A* RNAi lines were grown in a single climate chamber experiment for controlled day-length (16h light/8h dark; LD and 8h light/16 h dark; SD), at a temperature of 20°C day/18°C night), light regime of 300 μM/m^2^/s and relative humidity of 70%. Transgenic plants carrying the *StSP6A* RNAi (SP6Ai) construct were newly generated by *Agrobacterium* mediated transformation of *in vitro* internodes and 2 selected independent lines were used in the study. Lines were previously generated by Abelenda et al. (2019), however a new transformation was necessary as the lines used in Abelenda et al. (2019) were lost due to an infection in tissue culture. The plasmid generated there was used to retransform *S. andigena*. MS-20 tissue culture media with 2% (w/v) sucrose was used for *in vitro* vegetative propagation for three weeks. Tissue culture grown plantlets were planted to small pots (11×11 cm) and transferred to larger 3L pots 50 days after planting (DAP). Plants were initially kept in growth chambers for 4 weeks in LD conditions. After 4 weeks, the photoperiod was changed to SD to induce and synchronize tuber onset.

#### 2.1.2 Sampling of plant material and measurements

Destructive harvests of plants were performed to monitor above- and belowground growth. Starting from before the switch to short days (SD0), the harvest was initially performed weekly and later the frequency was lowered to ensure sampling of plants during senescence (Table 1). Plants were randomized in blocks within the climate chamber, from which at each time point five plants per independent line were randomly sampled to be destructively harvested. Error bars represent the standard deviation of five individually harvested plants, unless mentioned otherwise. For each genotype we thus started with a total of 50 plants. Stolon (number), tuber (number, fresh weight, dry weight, size distribution) and shoot (fresh weight, leaf area, stem length, stem diameter) were recorded for all plants (Table 2). For tuber dry weight, samples were dried at 70°C for 48 hours. For all timepoints, leaves, tubers and lower stem were harvested for sugar analysis. Soluble sugars were extracted and measured as described by Dinh et al. (2019). At SD8, the 1^st^ fully expanded leaves (∼5^th^ leaves from the top of the plant) were harvested for gene expression analysis at ZT3. RNA extraction was performed using the RNeasy Qiagen kit with on column DNase digestion using the RNase free DNase set and according to manufactur instructions. iScript cDNA Synthesis kit was used for cDNA synthesis starting from 1 μg of total RNA. Gene expression analysis was carried out in technical triplicates and the housekeeping gene *StElF3e* was used for normalization.

**Table 1.** Harvest timing.

| Harvest | SD0 | SD1 | SD2 | SD3 | SD4 | SD5 | SD7 | SD8 | SD10 | SD12 |
| --- | --- | --- | --- | --- | --- | --- | --- | --- | --- | --- |
| Days after planting (DAP) | 28 | 35 | 42 | 49 | 56 | 63 | 77 | 84 | 98 | 112 |

**Table 2.** Measured traits.

| *Aboveground* | *Belowground* |
| --- | --- |
| *Shoot Fresh Weight (sFW, g)* | *Tuber Fresh Weight (tFW, g)* |
| *Leaf Area (LA, cm<sup>2</sup>)* | *Tuber Dry Weight (tDW, g)* |
| *Stem Length (sL, cm)* | *Tuber Number (tN, -)* |
| *Stem Diameter (sD, mm)* | *Stolon Number (sN, -)* |

### 2.2 Data analysis and model descriptions

#### 2.2.1 Beta growth rate

To describe the growth curves of both leaf area and tuber fresh weight, we used the previously described beta growth function (Yin et al., 2003), which models determinate sigmoidal growth of plant organs. For tuber FW we use:

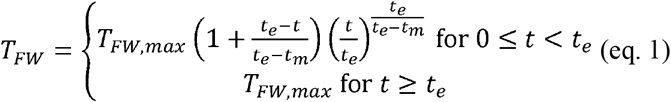

where T_FW_=0 at the start of growth (t=0) and T_FW_=T_FW,max_ at the end of the growth phase (t=t_e_), with maximum growth at t_m_.

Growth at specific time-points can be calculated with the equation below, which is the derivative of equation 1:

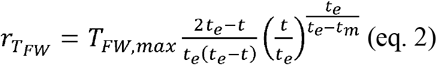

For the leaf area fit we extended the sigmoidal growth phase with an exponential decay after t_e_ to represent senescence:

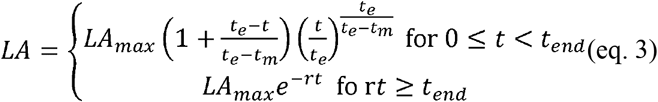

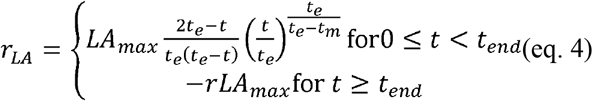

The tuber FW and leaf area of wildtype plants as well as each transgenic line were fitted individually using curve_fit from the scipy package. The ‘trf’ method was used as we set t_e_ boundaries per line to aid in obtaining a good leaf area fit for senescence start. The other two parameters (T_FW,max_/LA_max_ and t_m_) were not bound.

#### 2.2.2 Simulation of the stochastic onset of tubers

We created a simple model simulating tuber onset as a stochastic process with a tuberization probability. We assume that this probability consists of an SP6A dependent component (high when SP6A is present, zero when SP6A is absent) and an age dependent component (starting low, non-linearly increasing):

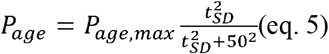

For SP6Ai lines the SP6A dependent probability always equals 0, for the wildtype plants we assume that this probability switches to a non-zero value after two weeks in SD conditions, with levels increasing from a baseline to a maximum over the next two weeks to model the time delay between leaf and tuber expression levels as reported by Navarro et al. (2011):

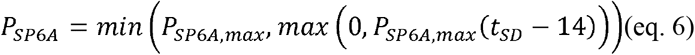

A predetermined number of stolons (“potential tubers”) is used, based on the average final number of tubers measured experimentally. We then simulate per day for each not yet tuberized stolon whether it will start tuberization or not. We do this by drawing a random number between 0 and 1, if this number is above the threshold (i.e. the tuberization probability at that day as determined by SP6A presence and plant age) a tuberization event occurs.

#### 2.2.3 ODE model of leaf and tuber growth

We developed a simple leaf area and tuber FW growth model where both LA and T_FW_ are assumed to grow proportional to leaf area, which would result in exponential growth of both leaves and tubers in absence of leaf senescence. T_FW_ was assumed to not decrease during the experiment, therefore we did not include a tuber decay rate in the model (equation 7). On the other hand, LA was observed to decrease due to senescence. To model this, we included a decay rate (d) for LA dynamics (equation 8). We investigated the effects of making this decay rate dependent on tuber size, and/or age and /or leaf area size (equation 9a). To model the potential effect of resource competition between leaves and tubers, we included a term describing increased senescence for larger tuber size (eq. 9b). This is based on the assumption that a larger tuber would be a stronger carbon sink. Developmental senescence was modeled dependent on the age of the plant (eq. 9c). Finally, we included a senescence inhibiting factor, which decreases the death rate for high leaf area, making sure that senescence due to resource competition is counteracted when carbon supply is abundant (eq. 9d). Terms 9b, 9c and 9d were applied in various combinations to determine which processes most accurately describe leaf senescence dynamics.

The equations were fitted to the LA and tuber FW data of WT and SP6Ai. Parameters were not constrained and are shared between the wildtype and transgenic lines, all differences in this model thus result from differences in tuberization timing. Parameter fitting was done using Grind.R by R.J de Boer (http://tbb.bio.uu.nl/rdb). Grind.R is an R script that functions as a wrapper for the deSolve and FME R packages (Soetaert et al., 2010).

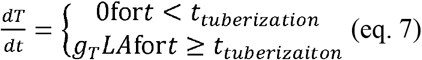

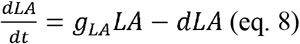

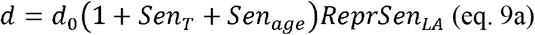

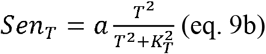

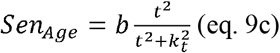

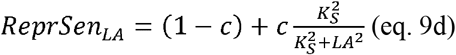

## 3. Results

### 3.1 Delayed canopy senescence concurs with continued apical growth in SP6Ai plants

*A priori*, it was expected that the knockdown of *StSP6A* would influence resource allocation and shoot versus tuber growth dynamics due to its effects on flowering and tuberization. Indeed, WT and SP6Ai plants did show large phenotypic differences in final stage shoot characteristics (Fig. 1A). The silencing of *StSP6A* was confirmed by qPCR analysis (Fig. 1C). Together, this confirms a strong phenotypic effect in *StSP6A* knockdown lines. In SP6Ai lines, expected to have enhanced flowering based on the flowering suppressive effects observed for *StSP6A* (Plantenga et al., 2019a), flowers emerged earlier and developed further (Fig. 1B).

**Figure 1.**
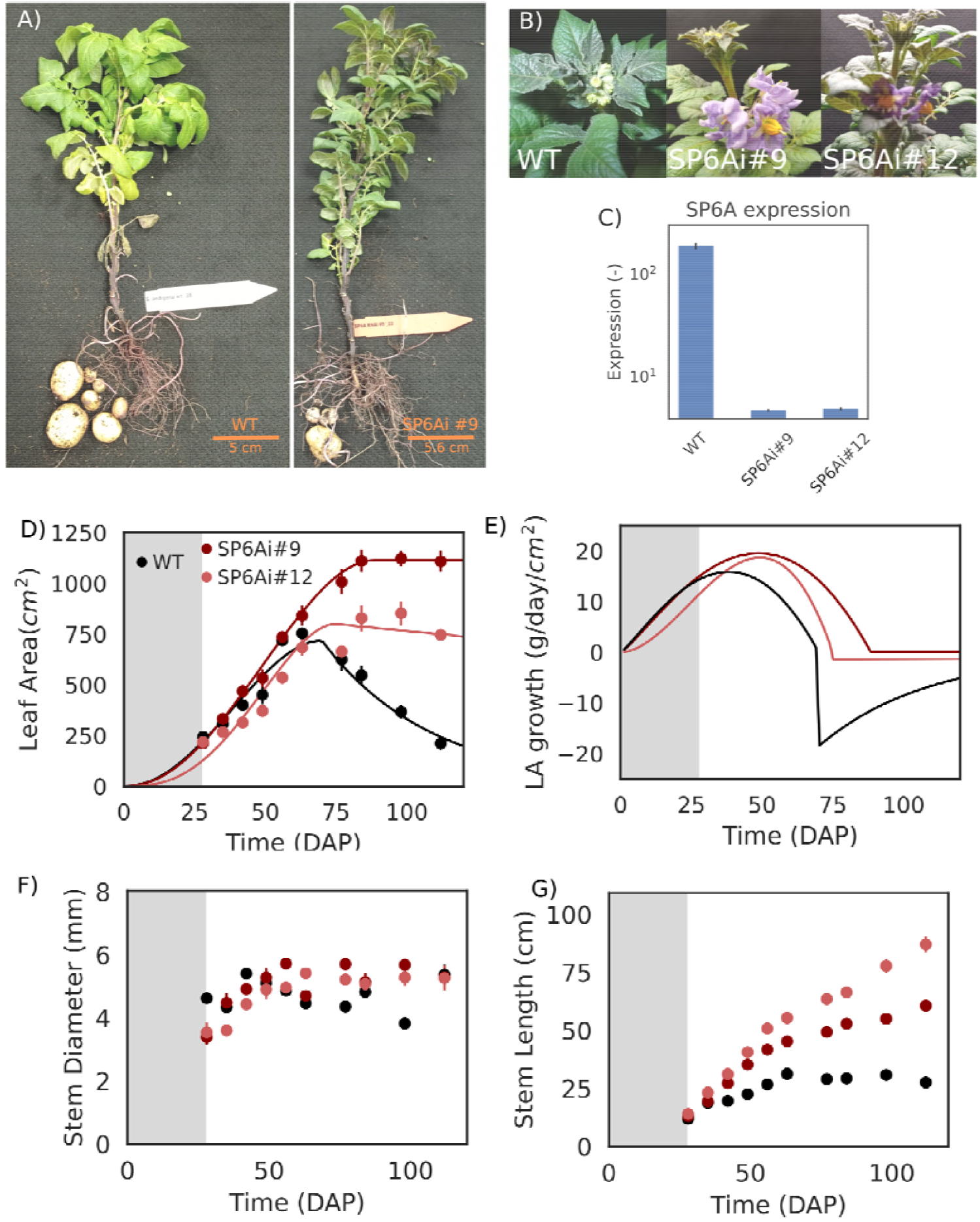
Aboveground characteristics and growth dynamics. **A)** Phenotypes of harvested plants at SD7. **B**) Flower and bud development at SD7. **C**) SP6A expression levels in WT and SP6Ai lines at SD8, error bars represent standard deviation of three independent measurements **D)** Leaf area over time, the gray area represents LD conditions, data was fitted using the beta growth function and an exponential decay function (see Methods). **E)** Leaf area growth rates over time **F)** Stem diameter as a function of time **G)** Stem Length as a function of time

During the early growth stages under long days (LD), no significant differences in leaf area were observed between the different plant lines (Fig. 1D grey area). Using a beta growth function, commonly used to fit determinate growth, to quantify growth rates, we also found no significant differences in leaf area growth rate (Fig. 1E grey area, see Methods). In the subsequent exponential growth phase, differences in leaf area and growth rate remained small. We did not observe an increased stem diameter for SP6Ai plants (Fig. 1F), opposed to recent observations of decreased stem diameter in *StSP6A* overexpression plants (Lehretz et al., 2021), suggesting a saturating dependence of radial growth on sucrose delivery. At later stages (>60 days after planting), stem length and leaf area started to show consistent and significant differences between the different lines (Fig. 1D, G). Looking at leaf area and growth rates, we observe that the increase in leaf area ended at 70-80 DAP for WT, followed by a decline in overall leaf area reflecting canopy senescence. Leaf area increase is prolonged in SP6Ai lines resulting in a larger final leaf area, with no substantial decline occurring within the timeframe of our experiment (∼120 days), representing a delay in canopy senescence of at least 4-6 weeks compared to the control plants. Specifically, apical elongation was observed to continue for SP6Ai lines (especially #12, Fig. 1G), resulting in an altered shoot morphology, with longer stems and leaf-loss restricted to the lower stem (Fig. 1A). As a result, senescence of older leaves was compensated by newly forming leaves, delaying overall leaf area decrease in SP6Ai lines (Fig. 1D). These results are consistent with previous experiments demonstrating an early suppression of apical growth in *StSP6A* overexpression lines (Lehretz et al., 2021). Summarizing, *StSP6A* transcriptional levels strongly correlate with overall canopy senescence timing.

### 3.2 Delayed tuberization in SP6Ai plants is mitigated by higher tuber growth rates

Next, we compared tuber growth dynamics (Fig 2), investigating tuber FW growth dynamics (Fig. 2A, B) tuber numbers (Fig. 2C), stolon numbers at SD1 (Fig. 2D) and overall final tuber FW (Fig. 2E) SP6Ai lines showed an approximately 2-week delayed tuberization onset, consistent with the 2- to-3-week delay observed in previous work (Navarro et al., 2011). Based on our beta growth function, in the three final harvests moments (SD8, 10, 12, i.e. 84, 98 and 112 DAP) higher tuber growth rates were estimated in SP6Ai lines compared to WT, with tuber growth continuing beyond the time when WT tuber growth halted (Fig. 2B). Overall, the combination of delayed tuberization onset and subsequent prolonged and elevated growth, resulted in an approximately similar (SP6Ai #12) or slightly higher (SP6Ai #8) final tuber FW (Fig. 2E) despite lower tuber numbers (Fig. 2C). In contrast, earlier results showed that SP6Ai plants had a reduced final tuber FW, similar to that of SWEETi plants (Abelenda et al., 2019). This inconsistency might be due to differences in final harvesting time, as we also observed lower FW for SP6Ai for the first 100DAP and only observed an increase compared to WT at the end of our longer experiments. Finally, tuber dry weight and FW show a consistent high correlation for all lines (Fig. 2F), indicating a constant dry matter content across lines. The increased tuber growth for SP6Ai is thus not merely caused by an increased water uptake but also represents an increased dry matter accumulation. Summarizing, under the later harvesting time applied here, the approximately 2-week delay in tuber onset can be approximately compensated by the 4–6-week delay in canopy senescence.

**Figure 2.**
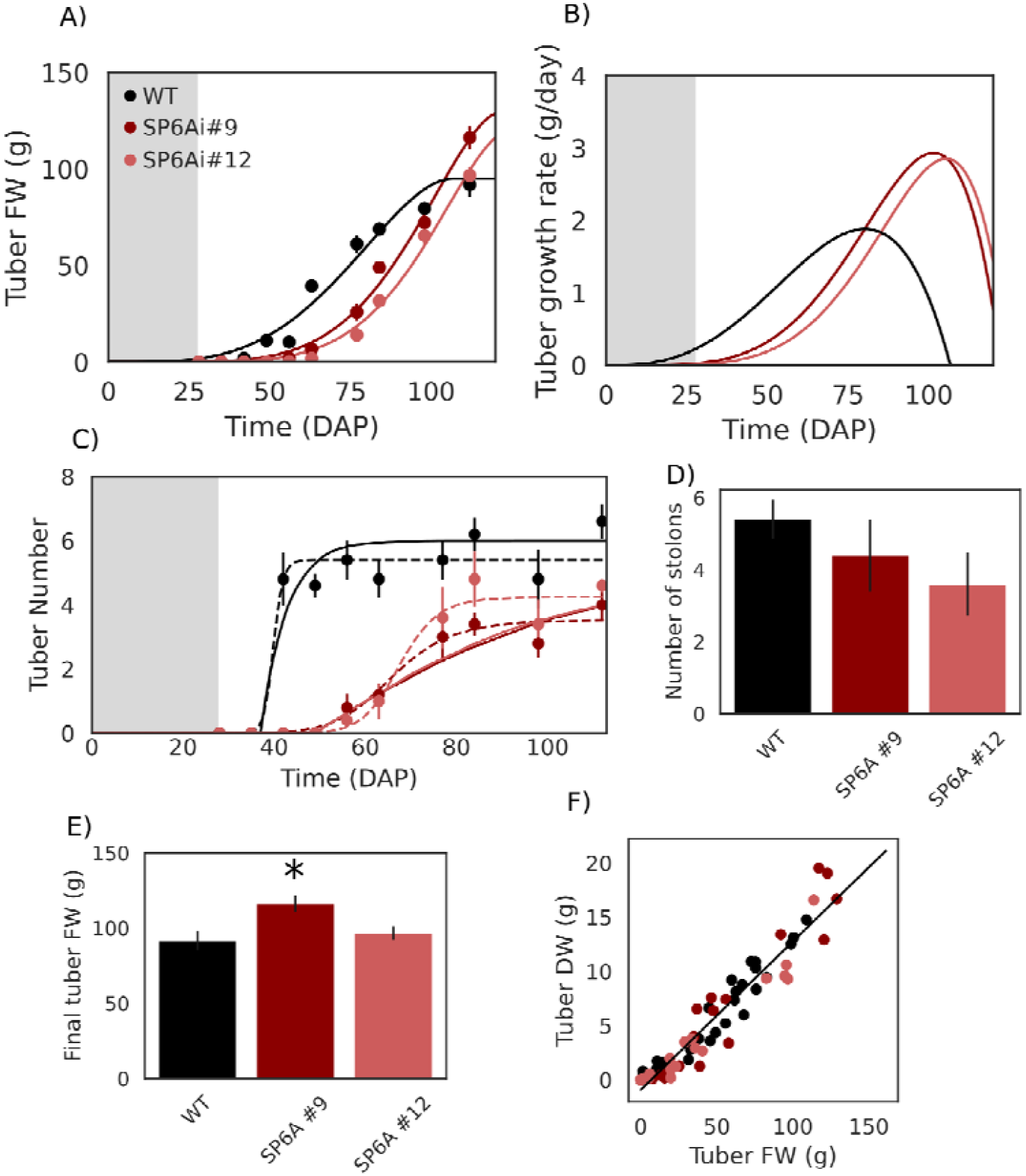
Tuber growth characteristics. **A)** Tuber FW over time, grey area represents LD conditions, data was fitted using the beta growth function **B)** Tuber growth rates over time **C)** Tuber number as a function of time. Dashed lines represent a sigmoidal fit, whereas solid lines represent the stochastic onset model **D**) Stolon number at SD1, just before the first visible signs of tuberization. **E)** Final harvest tuber FW. Stars represent a significant difference with WT (students t-test, p<0.01) **F)** Correlation between tuber FW and DW (r=0.95, p<0.005)

### 3.3 StSP6A knockdown causes gradual tuberization

In addition to differences in leaf and tuber growth dynamics and final tuber FW, we also observed differences in tuber number dynamics (Fig. 2C) and number of stolons present before tuberization (SD1, Fig 2D). The increase in tuber number for SP6Ai lines was significantly more gradual (dashed lines, Fig. 2C) and reached a lower final number of tubers. This is consistent with the increased tuber numbers observed in *StSP6A* overexpression reported by Lehretz et al. (2021). In all lines the number of initial non-swelling stolons matches the number of final tubers formed (Fig. 2C) indicating that stolon formation is not a limitation factor, leading to the more gradual formation of tubers in SP6Ai lines. Instead, the more gradual increase in tuber numbers suggests that *StSP6A* may play an important role in tuberization synchronization as well as tuber onset.

To investigate this, we created a simple phenomenological model simulating tuber onset as a stochastic process with a tuberization probability. We assumed that this probability consists of the sum of an *StSP6A* dependent component (high when *StSP6A* is present, zero when *StSP6A* is absent) and an age dependent component (starting low, non-linearly increasing with time) (see Methods). We start with the predetermined number of stolons as observed in the experiment. It is then determined per day for each stolon whether it will start tuberization or not. This model was used to estimate the *StSP6A* dependent and independent probabilities based on the experimental data. For the SP6Ai line we assumed that the *StSP6A* dependent component of the tuberization probability always equals zero. For the other lines we assume that this component switches to a non-zero value after two weeks in SD conditions, with levels increasing from a baseline to a maximum over the next two weeks (see Methods), to model the time delay between leaf and tuber *StSP6A* mRNA expression levels as reported by Navarro et al. (2011). Fitting this simple model indeed enables us to closely mimic WT and SP6Ai tuberization dynamics (Fig. 2C, solid lines) and results in a fitted per-day *StSP6A*-dependent tuberization probability of 15-20% and a *StSP6A*-independent tuberization chance increasing from 1 to 3.5% per day over time. This suggests that *StSP6A* acts as a strong signal for synchronization of tuberization. Importantly, while we do not explicitly consider other regulatory factors such as *StBEL5* or *StSP3D*, our model results indicate that these factors should either gradually increase over time, having similar effects as age, or instead have only a small effect on daily tuberization chances as otherwise SP6Ai results can not be fitted.

### 3.4 Gradual tuberization correlates with increased variance in tuber size distribution

Tuber size distribution is an important agronomic trait, and variability in tuber size has been attributed to heterogeneity in seed tuber storage and growth conditions, stolon length and hierarchy as well as differences in tuberization dynamics (Aliche et al., 2019; Ospina et al., 2021; Struik, 2007). Therefore, we investigated whether the observed differences in tuberization dynamics correlate with differences in tuber size distribution. While we have not measured all individual tuber weights and hence cannot calculate tuber variance directly, we estimated the variability in tuber size distribution from the available data by taking the weight ratio between the largest tuber (g_M_) and the average weight of the other tubers (g [_rest_):

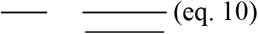

The higher this ratio, the larger the difference between the largest tuber and the average of the rest is, enabling us to use it as a proxy for variation in tuber size distribution. We clearly see that, at the final harvest for SP6Ai, the largest tuber is significantly larger (Fig. 3A), while the average of the other tubers is lower (Fig. 3B), resulting in a higher ratio (Fig. 3C). This suggests that the delayed and gradual onset of the SP6Ai tubers correlates with an increase in variation in tuber size distribution under pot conditions.

**Figure 3.**
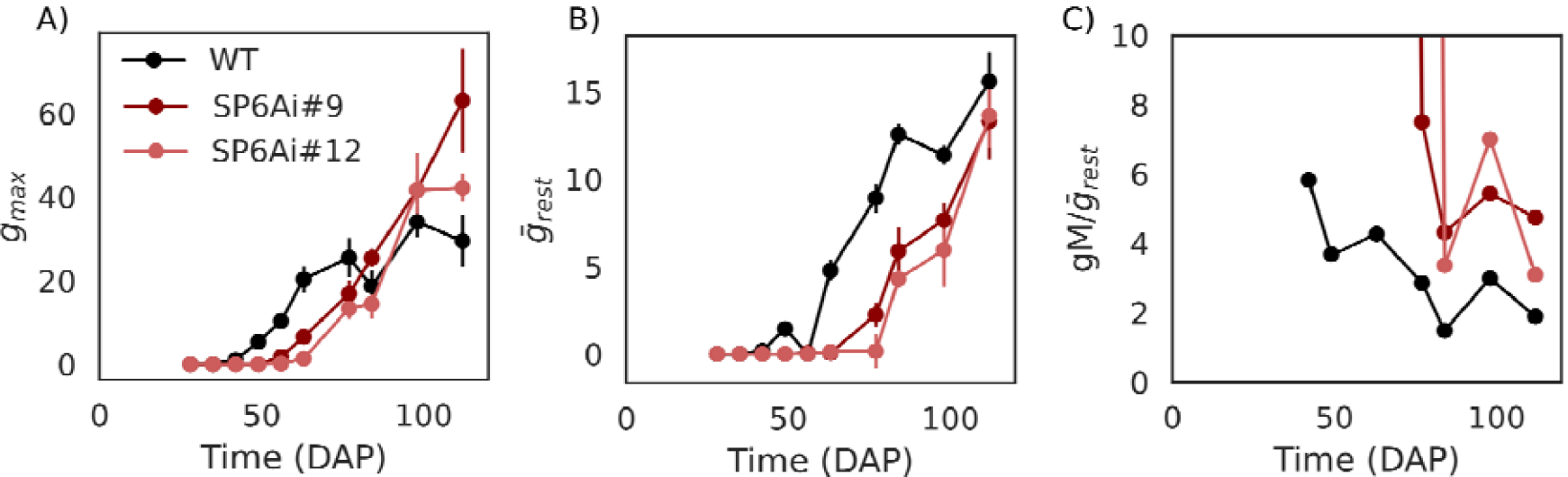
A proxy for tuber size distribution variation. **A)** Largest tuber dynamics **B)** Average size of all tubers except the largest tuber **C)** The ratio of the largest tuber over the average of the other tubers, the larger this ratio is, the larger the variation in tuber size. The ratio during early tuberization in the SP6Ai was >100 due to gradual tuber onset. For readability the y-axis was set from 0 to 10.

### 3.5 A negative correlation between tuber and leaf growth suggests a role for resource competition

Leaf area, tuber FW and tuber number dynamics described above demonstrated a strong correlation between timing of the onset of canopy senescence and the onset and halting of growth of tuber FW. To investigate potential mechanisms underlying these observations, we first investigated the interdependence between leaf area and tuber and leaf growth dynamics. Tuber FW did not correlate strongly with leaf area (Fig. 4A, Pearson’s r=0.17, p<0.094), yet tuber growth rate does strongly correlate with leaf area (Fig. 4B, r=0.79, p<0.005), consistent with leaf area determining photosynthetic output and thereby growth potential. This correlation furthermore supports our hypothesis that delayed canopy senescence and resulting larger cumulative leaf area causes the enhanced and prolonged tuber growth observed in SP6Ai lines.

**Figure 4.**
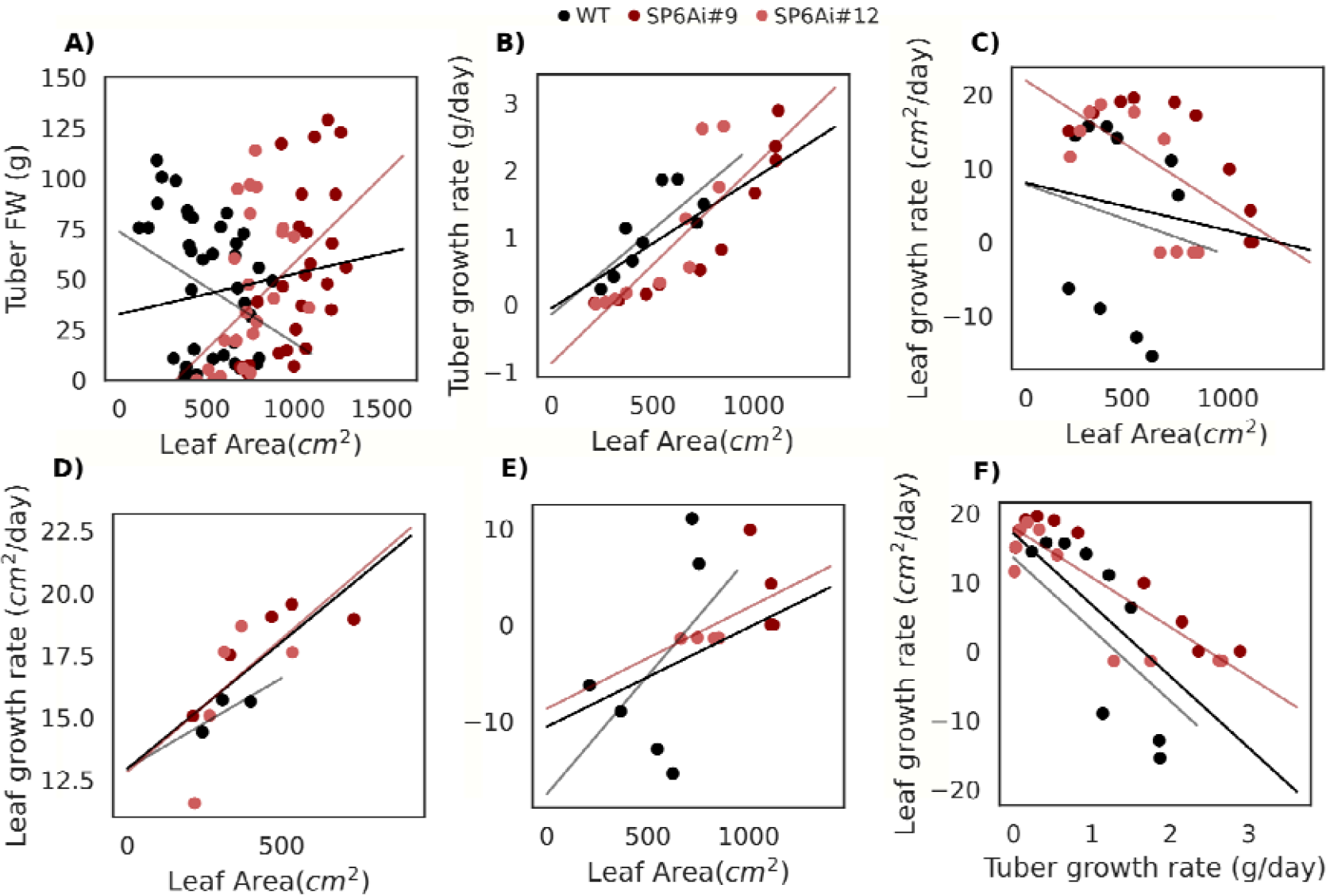
Correlations between Leaf Area, tuber FW and tuber and leaf growth rates. **A)** Leaf area vs. Tuber FW. Black line depicts a regression for all lines combined. Colored lines depict a regression for their transgenic line respectively. **B)** Leaf area plotted against the tuber growth rate Leaf area plotted against the leaf growth rate **D)** Leaf area plotted against the leaf growth rate pre tuberization (<SD2 for WT and <SD4 for SP6Ai lines), **E)** Leaf area plotted against the leaf growth rate post tuberization (<SD2 for WT and <SD4 for SP6Ai lines). The dashed line represents the post-tuberization senescence trajectory. **F)** Tuber growth rate plotted against the leaf growth rate

Next, we investigated the relation between leaf growth rate and leaf area (Fig. 4C), for which we found a weak, non-significant, negative correlation (r=-0.16, p=0.18). Given the clear distinction between an overall leaf area growth and senescence phase (Fig. 1D-E), we reinvestigated this relation while separating leaf growth rates occurring pre- or post-tuberization. Pre-tuberization leaf growth rate and leaf area displayed a positive correlation (Fig. 4D, r=0.68, p<0.005), while for post-tuberization only a weak correlation was observed (Fig. 4E, r=0.23, p=0.015). Thus, in absence of tubers, leaf area drives leaf growth as expected, yet post tuberization leaf area only drives tuber growth while simultaneously declining due to senescence. In agreement with the correlated timing of tuberization and canopy senescence onset, we observe a strong negative correlation between tuber and leaf growth rates (Figure 4F, r= -0.79, p<0.005). Overall, these correlations suggest a potential role for resource competition in canopy senescence and its timing.

### 3.6 Sucrose concentrations do not clearly affect growth dynamics after tuberization

While our data demonstrate a strongly correlated timing of tuberization onset and canopy senescence, a major question remains whether this is caused by a shared, possibly *StSP6A* dependent regulation or rather arises from tuberization induced resource competition inducing/enhancing senescence, or from a combination. Indeed, senescence has been suggested to be promoted by both elevated as well as lowered sucrose levels, independent of developmental age (Rankenberg et al., 2021; Wingler, 2018). To investigate the hypothesis that resource competition enhances senescence, we measured sucrose levels in leaves, stem, and tuber over the course of development in the wildtype and SP6Ai lines (Fig. 5A-C). The sucrose data showed elevated levels in the stem for SP6Ai lines, confirming earlier findings on the role of *StSP6A* in reducing *StSWEET* mediated sucrose export (Abelenda et al., 2019). We did not observe a decrease in leaf sugar levels after tuberization onset in any of the lines to support our hypothesis of competition mediated low sucrose levels in leaves enhancing senescence.

**Figure 5.**
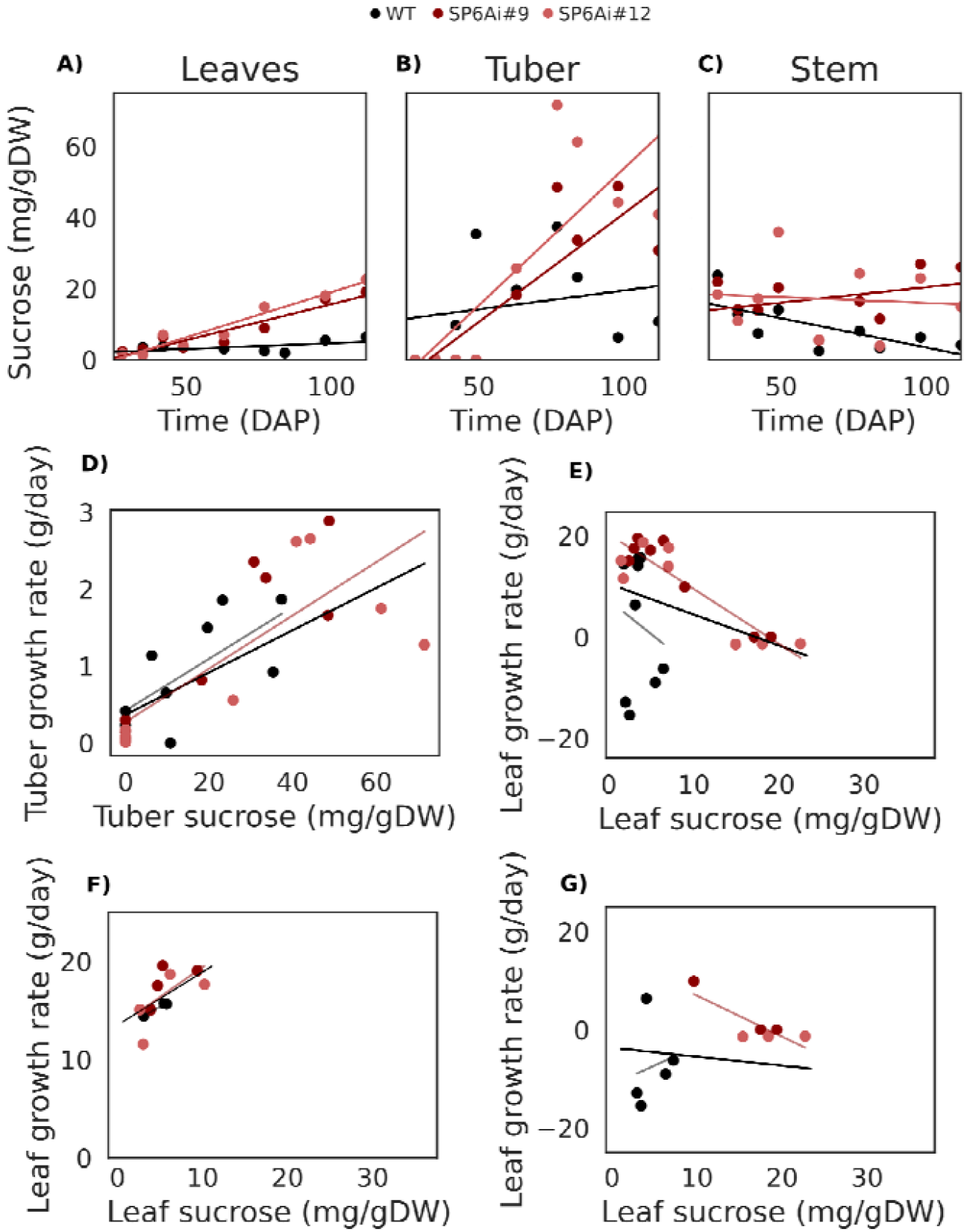
Sucrose dynamics and its relation to organ size and growth. **A)** Sugar concentration dynamics in tubers, a linear regression was done to visualize the general time dynamics of sucrose levels. **B)** Sugar concentrations dynamics in leaves **C)** Sugar concentration in whole stem samples **D)** Tuber sucrose vs. tuber growth rate **E)** Leaf sucrose vs. Leaf growth rate **F)** Leaf sucrose vs. Leaf growth rate pre-tuberization **G)** Leaf sucrose vs. Leaf growth rate post-tuberization

To further probe for signals of resource, we correlated organ sugar levels with organ growth (Fig. 5D-G). We find that tuber growth rate is positively correlated with tuber sucrose levels (r=0.72, p<0.005, Fig. 5D) for the entire duration of the experiment and for all lines. Leaf sucrose levels are negatively (r=-0.55, p<0.005, Fig. 5E) correlated with leaf growth rate, yet when again splitting pre- and post-tuberization leaf growth rates, we observe a positive correlation (r=0.63, p<0.005, Fig. 5F) pre-tuberization. Post-tuberization there is again no clear correlation present (r=--0.09, p=0.94, Fig. 5F). Instead, growth/senescence rates are relatively stable within the transgenic lines for a range of sucrose concentrations. Thus, the sucrose data did neither indicate that tuberization lowers leaf sucrose levels (Fig. 5A) nor that sucrose levels strongly determine leaf growth rate (Fig. 5E-G).

### 3.7 A simple leaf and tuber growth model suggests a role for resource competition in delayed senescence

While our sugar data do not provide indications for resource competition between tubers and leaves, this may be due to the limited spatial resolution of our sugar measurements. To further investigate the mechanism underlying the correlated timing of tuberization and canopy senescence onset we therefore developed a simple leaf and tuber growth model (see Methods for details). In the model, we assumed that leaf and tuber growth rate are proportional to leaf area, consistent with our observations (Fig. 4B, D), and additionally incorporated leaf senescence. Importantly, our data demonstrate that canopy senescence in SP6Ai lines is further delayed than tuberization onset, suggesting a non-trivial interrelation, potentially involving sugar status and/or other signaling pathways. Therefore, for canopy senescence we investigated the potential impact of tuber size and plant age promoting, and total leaf area inhibiting canopy senescence.

As a baseline, we performed a simulation with a constant leaf death rate. The best fit (Fig. 6, dotted lines) did neither qualitatively nor quantitatively represent the experimental data well. Specifically, no transition from leaf area expansion (leaf growth rate exceeding leaf turnover) to overall shoot senescence (leaf turnover exceeding leaf growth) could be achieved, due to the constant leaf turnover rate. An additional tuber dependent leaf death rate was therefore added, where increasing tuber FW leads to a higher leaf death rate (Fig. 6, dashed lines). In the best fit, no substantial overall shoot senescence occurred, instead leaf area became constant. This can be understood from the interplay between tuber size and leaf area, with tuber size negatively impacting leaf area, yet tuber growth positively depending on leaf area. As a result, a balance is reached when leaf growth and turnover are equal, resulting in constant tuber growth. As WT and SP6Ai lines displayed different tuber growth dynamics, this transition to constant leaf area and linear tuber growth occurred at different time points.

**Figure 6.**
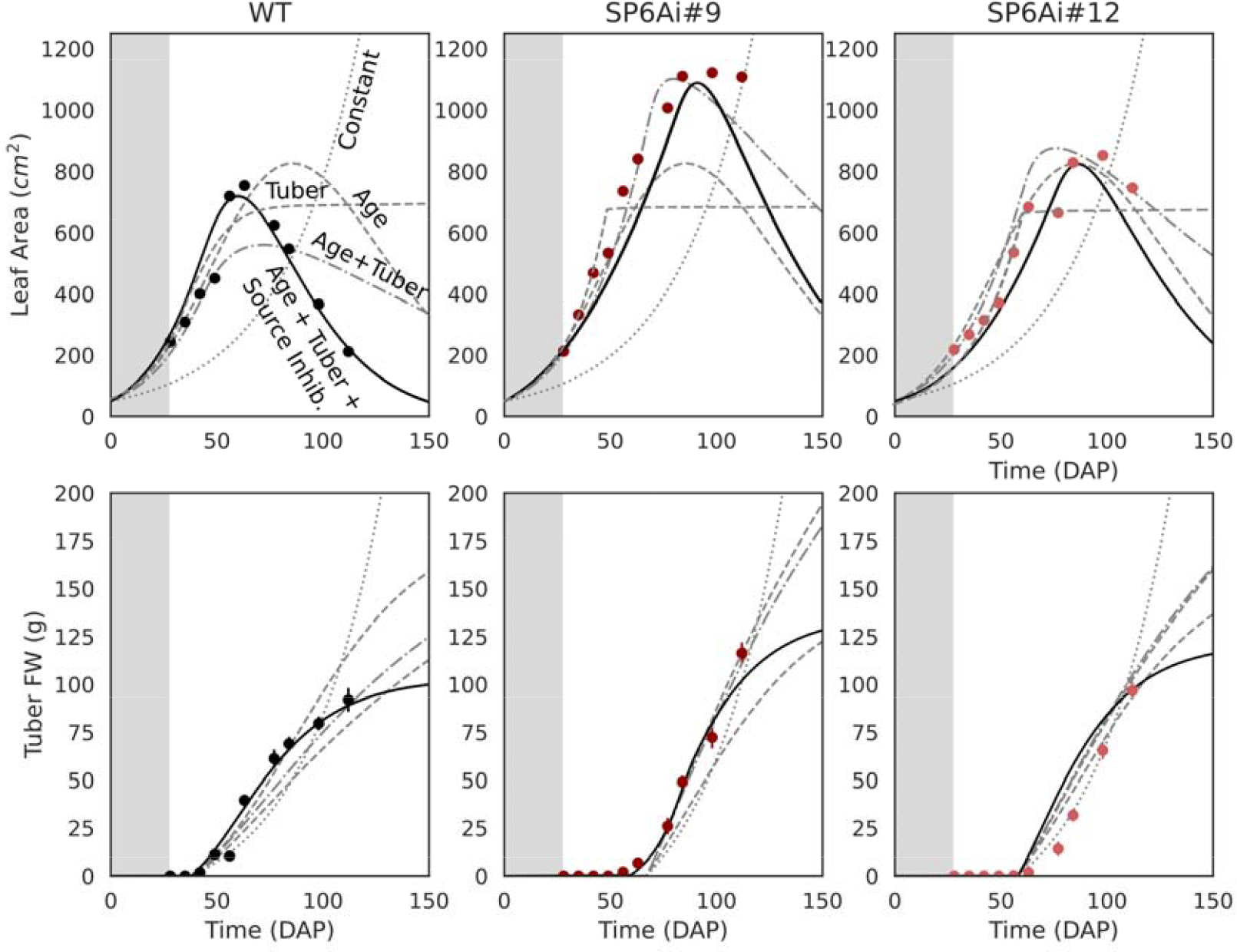
Leaf Area and Tuber FW fitted with the growth model. Best fit (solid lines) to the leaf area and tuber FW data (dots) using the growth model in solid black lines, for WT and SP6Ai. The other model variants are depicted with different line types, i.e., Constant (dotted), Age (dashed), Tuber (large dashed), Tuber+Age (dash-dotted).

Besides senescence due to resource competition, plant age (i.e. DAP) can be a prominent cause for canopy senescence (Wingler et al., 2018). We therefore included a plant age dependent death rate, adding this to the baseline model with only a constant death rate (Fig. 6, large dashed lines). This led to a best fit where overall shoot senescence started at around 75 DAP for all plant lines. This consistent senescence timing is too late for WT and too soon for SP6Ai lines and arises from the inability of this model to discriminate between the different plant lines. We next combined age and tuber size dependent models (Fig. 6, dash-dotted line), enabling the model to discern plant lines based on differences in tuber size development dynamics. Nevertheless, the best fit was still dominated by plant age resulting in a similar senescence timing for all transgenic lines.

Tuberization onset is delayed by only 2 weeks between SP6Ai and WT, whereas canopy senescence is delayed considerably longer. This may explain why tuberization is insufficient to explain the differences in canopy senescence timing. Differences in leaf area dynamics and maximum achieved canopy sizes affect overall sucrose source strength and may thereby impact the severity of resource competition between tubers and leaves. We therefore hypothesized that differences in tuberization onset may lead by themselves to minor differences, which through affecting source strength, become amplified to larger differences in senescence. We included this hypothesis through the addition of a term that inhibits senescence based on the size of the canopy (source inhibition, Fig. 6, solid lines). We found that only when including a combination of age, tuber and canopy size effects on leaf senescence rate, we could reproduce our experimentally observed leaf area and tuber FW dynamics.

While this model cannot proof causality, by varying the threshold level at which tuber size enhances senescence, we can investigate whether tuberization onset and thus induction by a shared signal (low tuber size threshold: K_T_ < Tuber FW, so rapidly saturating) or rather growth of tubers and thus resource competition (high tuber size threshold K_T_ > Tuber FW, so gradually increasing) better explains canopy senescence dynamics. The best fit to the experimental data was found to be K_T_ 117gFW, which is on the higher end of observed tuber FW (see Figure 2A), suggesting that with regards to the tuberization dependent part of senescence control resource competition rather than a shared signal underlies the correlated timing of tuberization and canopy senescence.

## 4. Discussion

In this study we performed a temporally resolved climate chamber experiment in combination with simple models to investigate the impact of the presence and timing of different sink organs, the temporal dynamics of source organs, and the interplay between sink and source organs on final tuber FW.

Consistent with earlier findings, we observed a 2-week delay in tuberization in SP6Ai lines (Navarro et al., 2011). Additionally, tuberization appeared more gradual in these lines, and our experimental data and modeling suggest an important role for *StSP6A* in synchronization of tuberization. Still, the molecular mechanism underlying this synchronization remains to be investigated and could be either through *StSP6A* functioning in switching on tuberization genes, or through *StSP6A* reducing SWEET-mediated sucrose loss with the enhanced sucrose availability dampening inter tuber differences, or a combination of such processes.

Our data further suggests that this gradual tuberization correlates with a larger variation in tuber sizes (Fig. 3). While differences in tuber onset timing likely play a role in generating this variation in size differences, knockdown of *StSP6A* may have also unmasked other differences. To further substantiate the role of *StSP6A* in tuber onset synchronization and size homogenization, future studies applying rhizotron based imaging of the growth dynamics of individual tubers in WT and *StSP6A* knockdown lines may help decompose the effects of differences in stolon length, width, and hierarchy as well as tuberization timing on final tuber size.

In line with the generally observed ‘post-tuberization senescence’, we observed a strong correlation between tuberization and canopy senescence dynamics, with SP6Ai lines displaying both the latest tuberization and senescence. Interestingly, the combined early and late time points of final harvesting in our study enabled us to observe that the delay in senescence (4-6 weeks) was substantially more pronounced than the delay in tuberization onset (2 weeks), indicating this is not a simple, homogeneous delay in plant development. Furthermore, we observed that final tuber FW in the two SP6Ai lines was similar compared to WT, whereas in earlier time points SP6Ai tuber FW is (far) below WT, as was also observed previously (Navarro et al., 2011; Abelenda et al., 2019). This suggests that delayed and more gradual tuberization can be compensated by even more delayed canopy senescence, underlining the importance of considering both sink and source timing dynamics. Indeed, Struik & Wiersema, (1999) already stated that “Fastest overall development is not necessarily associated with the highest yields”, later confirmed by Aliche et al., (2018), and further supported by our data.

Therefore, we sought to investigate whether the non-trivial correlation in tuberization and canopy senescence timing primarily arises from either a shared inductive signal, a signal downstream of tuberization inducing senescence or rather that there is also a significant role for resource competition between tubers and leaves in canopy senescence timing. We observed clear positive correlations between leaf area and tuber growth rate in all transgenic lines, as well as tuber sucrose levels and tuber growth rate, supporting a prominent role for source size in tuber growth. In contrast, while we observed an inverse relation between leaf and tuber growth rates suggestive of resource competition, neither a decrease in leaf sucrose levels post tuberization nor a clear correlation between leaf sucrose levels and growth rate was found either pre- or post-tuberization. Thus, our sucrose data did not provide clear indications for resource competition between tubers and leaves.

This could either imply that competition for sugar does not play a key role in the coordinated timing of tuberization and canopy senescence or rather indicate that the coarse-grained spatial resolution of the applied sucrose measurements, measuring overall sucrose levels irrespective of tissue in only a single leaf per plant, is insufficient to resolve this competition. As an alternative means to investigate potential causal relations between leaf and tuber growth dynamics we developed a simple leaf and tuber growth model, which we fitted to data for WT and SP6Ai lines. Good model fits for canopy dynamics could only be obtained when senescence was not only age dependent but also promoted through gradually increasing tuber size, rather than switching to a higher level beyond a certain tuberization level. Additionally, model fits improved by incorporating that leaf area, and thus source strength, delayed senescence. Combined this model suggests that resource competition rather than a shared signal underlies the correlated timing of tuberization and canopy senescence, and that a complex dependence of senescence on the strength of competing sinks and overall source strength may explain how smaller differences in tuberization timing can become translated into larger differences in canopy senescence timing. Clearly, these model findings neither proof causality, nor that sucrose is the predominant factor next to age controlling leaf senescence. One major trigger for early senescence not considered in this study is the absence of sufficient nitrogen, which leads to degradation of chlorophyll to remobilize nitrogen (Gan & Amasino, 1997; Kim et al., 2009; Qiu et al., 2015). Modeling work done in other crop species, such as wheat, have shown that the carbon/nitrogen (CN)-balance proved to be pivotal in understanding the role of protein turnover and leaf senescence in understanding grain yield (Barillot et al., 2016). An interesting avenue for further investigation could be to use models to investigate to what extent tuberization onset impacts sucrose allocation to roots, thereby impacting their nitrate uptake efficiency and growth, thus indirectly enhancing shoot senescence.

Summarizing, our climate chamber experiments and models strongly suggest a role for resource competition in the correlated timing of tuber onset and canopy senescence as well as a leading role for *StSP6A* in tuber onset synchronization and tuber size uniformity. Future studies measuring gene expression, sucrose and nitrate dynamics in different leaves and tissues, and measuring not only leaf and tuber but also root growth dynamics are needed to pinpoint the exact role of resource competition in canopy senescence timing. Likewise, more detailed measurements are needed to reveal the molecular mechanisms through which StSP6A synchronizes tuberization. Finally, a major open issue is to what extent our results can be generalized from the climate chamber to the field.

## Acknowledgements

We thank Rutger Hermsen for input on the statistics of tuber size variability and Samjhana Khanal for assisting in the harvests.

## Authorship

BH performed growth experiments and sugar analysis and developed, implemented, analyzed the models and data. SB performed and designed the growth experiments, qPCR and sugar analysis. CB conceived the project. KT conceived the project and analyzed the data and models. All authors contributed to writing the manuscript and approved the submitted version.

## Data availability

Generated data is included in the supplementary material.

